# *Limacus flavus* yellow slug: bioactive molecules in the mucus

**DOI:** 10.1101/2021.05.06.442857

**Authors:** Patricia Yumi Hayashida, Pedro Ismael da Silva Júnior

## Abstract

**Background:** Snails and slugs were used as a treatment for many health problems therefore ancient times. Since the antimicrobial resistance became a major global thread, antimicrobial peptides have been considered as a potential source for development of new drugs, especially for drug-resistant bacteria. Nowadays reports confirm that the mucous secretions have antimicrobial, antiviral and antifungal properties.

**Methods:** The present study has the objective to characterize and evaluate antimicrobial peptides of *Limacus flavus* mucus. The mucus was obtained by thermal shock and submitted to RP-HPLC. Fractions were used to perform the antimicrobial activity and hemolytic assays, electrophoresis (SDS-Page Gel) and submitted to mass spectrometry (LC-MS / MS). Identification and characterization was performed by PeaksX+ software. The physicochemical parameters were evaluated with bioinformatics tools, which predicted water solubility, iso-electric point, charge net and its primary structure.

**Results:** Three fractions were isolated from the mucus of *L. flavus* and presented antifungal and antibacterial activity. The mucus showed greater inhibition for filamentous fungi (*Aspergillus niger*), yeast (*Cryptococcus neoformans*), Gram positive bacteria (*Bacillus subtilis, Micrococcus luteus*) and Gram negative bacteria (*Enterobacter cloacae*). These fractions also did not show hemolytic activity for human blood cells (erythrocytes). Fraction’s sequences were identified and presents Mw <3kDa, WLGH, DLQW, YLRW, respectively.

**Conclusion:** This study revealed three antimicrobial peptides of *L. flavus* mucus with a wide range of antimicrobial activity and its physic-chemical characterization.

## 1. INTRODUCTION

Antimicrobial resistance (AMR) is considered one of the major crises in the public health system worldwide, where attempts are made to suppress bacteria resistance to drugs [1]. For the past 30 years, antimicrobial peptides (AMPs) have been considered as a potential source of development of new antimicrobial drugs, specifically for multi-resistant-drugs. Although, evolutionarily, microorganisms, whether bacteria, viruses, fungi or parasites, suffer selective antimicrobial pressure, leading them to develop and acquire a natural resistance to existing drugs [2]. Thus, due to inefficient control of infections, the geographical movement of infected people and animals and environmental contamination increase resistance locally and globally [3].

AMPs are mostly amphipathic and cationic molecules, structurally composed of 5 to 100 amino acid residues [4,5]. They are produced by both vertebrates and invertebrates and plants [4]. They have a broad spectrum of activity against bacteria, fungi, protozoa and viruses [4,6], according to their physical-chemical structure, which includes their size, electrical charge, amphipathic structure, hydrophobicity and the mode of action [4,6]. Also, AMPs can have more than one biological response exhibiting a range of different activities, such as antiparasitic, antitumor, antiobesity, stimulates cell proliferation, angiogenesis and Vasculongenian properties, promotes wound healing and inhibits the inflammatory response, in addition to acting directly in the body’s antimicrobial defense. As it presents a variety of responses, AMPs have been increasingly studied; both to complement information about their functional properties and for pharmacological industrial interest [7].

Mucus is a complex mixture of products of great variety qualitatively and quantitatively, which forms a mechanical and biological barrier on an epithelial surface. This fluid is considered biphasic because it has an aqueous phase and a gel phase [8]. Being composed of a high water content and presents molecules of high molecular weight, in addition to gel-forming molecules [9]. For example, glycoproteins (mucins)[9], proteins, proteoglycans and lipids [10]. Its composition is similar between the most derived vertebrates and invertebrates [11].

Mucus is considered a highly versatile material, being produced for several functions: breathing [9,12], ionic and osmotic regulation [9,12], reproduction (gamete exchange) [9,12], nest production [9], excretion [9], communication (hormones and smells) [9,12], feeding [9,12], locomotion [13], adhesion to surfaces [12], hydration [10,14–16], mechanical protection and disease resistance [9,10,12,17,18].

Some of the research related to mucus is of snails and slugs, which have been in human health since ancient times, with their importance in folk and traditional medicine [19]. Snail mucus is well described in the literature, with biological and chemical properties in detail [19]; however, this does not occur in terrestrial slugs.

In slugs, mucus contains lectins [20], mucopolysaccharide and glycoprotein [20]. In the *Arion ater* slug, Cottrell et al. [21] found glycosaminaglycan with high concentrations of galactosamine and galactose. In addition, Otsuka-Fuchino et al. [22] reported in the *Achatina fulica* mucus a glycoprotein, achatin, which has antimicrobial activity against Gram-positive and Gram-negative bacteria. Some authors such as Iguchi et al., [14], Kubota et al. [15], Otsuka-Fuchino et al. [22], Toledo-Piza [17,18] and Araujo [23] demonstrated that slug mucus has antimicrobial peptide.

One of the most recent researches was carried out by the group of Li et al [24] where it was possible to carry out the transcriptome of the total body of the slug *Limacus flavus* with the next generation sequencing technique. The data resemble similarity with *Aplysia californica, Lottia gigantean* and *Crassostrea gigas* sequences. Furthermore, AMPs and protein-like were identified, such as lysozymes (antibacterial activity), defensins (antibacterial, antifungal and antiviral activity), thaumatin-like protein (antifungal activity), peritrophin (antibacterial activity), cystatin (antibacterial, antifungal and antiviral activity) and others fragments of AMPs which have antimicrobial activity and wound healing functions.

## 2. MATERIALS AND METHOD

### 2.1 YELLOW SLUG

Adult specimens of *L. flavus* were collected in São Paulo/SP, Brazil, at Vila Sônia neighborhood (23°59’54”S 46°73’44”W). They were stored alive in plastic compartments for transportation to the Laboratory of Applied Toxinology (LETA), at Butantan Institute, and stored in plastic boxes (33.0 x 21.8 x 10.3 cm) with pierced lids at 20-25°C. They were fed two times a week, with mice’s ration or lettuce; and in a plastic container with wet cotton. The cleansing of the materials was made three times a week.

### 2.2 MUCUS EXTRACTION

The animals were kept for three days without food before mucus extraction. They were submitted to thermal shock, which consist the submersion of each one in cold ultrapure water. Subsequently, the mucus was obtained by scraping the body with the help of a wooden spatula and stored in a sterile container (Figure 1). The material was subjected to lyophilization and stored at −80 °C.

**Figure 1.**
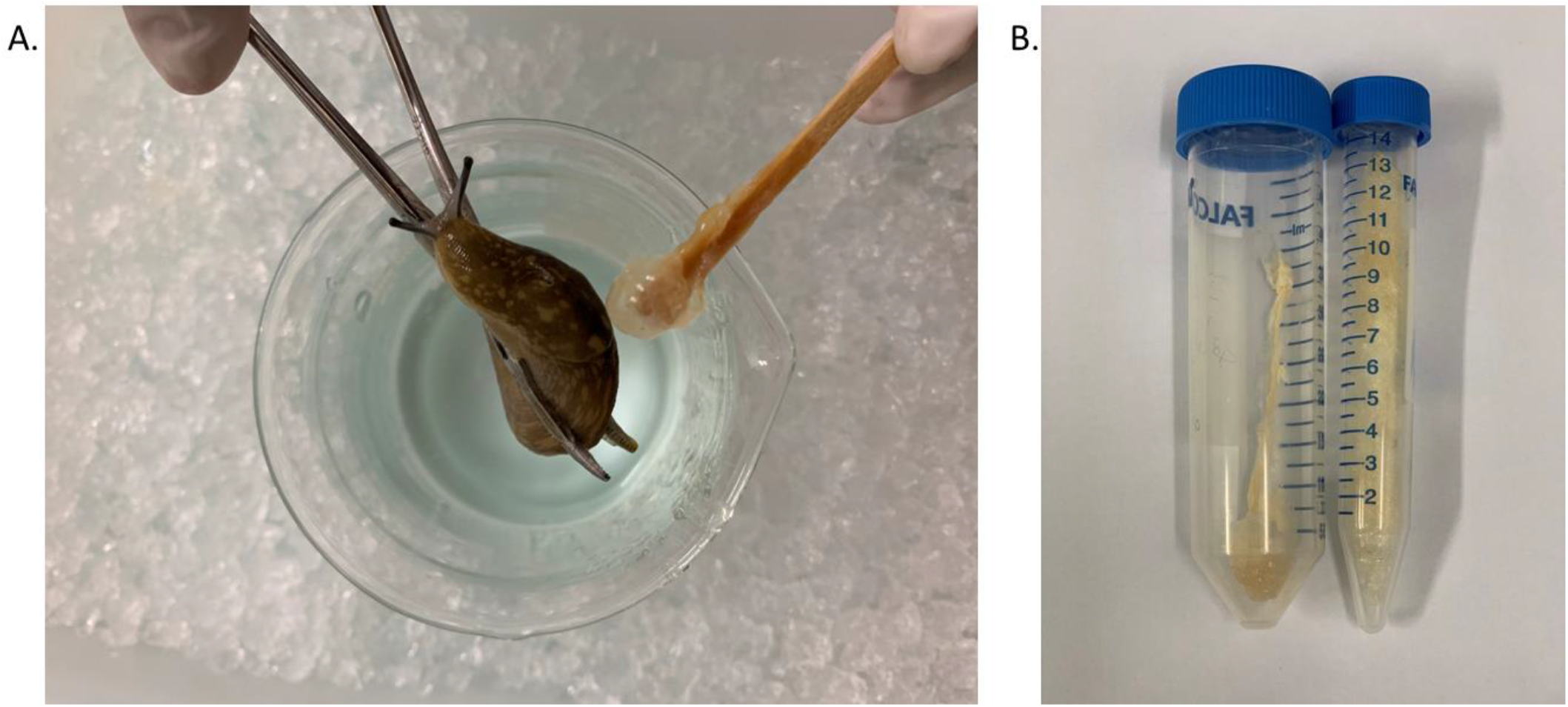
*Limacus flavus* slug (A) Mucus extraction from the *Limacus flavus* slug by thermal shock and body scraping. (B) Lyophilized raw mucus stored in a conical centrifuge tube of 15.0 mL and 50.0 mL.

### 2.3 REVERSE-PHASE HIGH-PERFORMANCE LIQUID CHROMATOGRAPHY (RP-HPLC)

Mucus was homogenized with 5.0 mL 10% DMSO for 2 min on a shaker, centrifuged (Centrifuge5804R Eppendorf ^®^ Instruments, Inc.) for 5 min, at 14000 x g at 4 °C. RP-HPLC purification was performed with a Shim-pack XR-ODS preparative column (5 μm; 20 mm; 250 mm, Shimadzu^®^), coupled with a preparative Shimadzu^®^ system, with a gradient of 0% to 80% acetonitrile (0.1% TFA), over 60 min, with flow rate of 8 ml / min. Fractions were collected manually and the absorbance monitored at 225 nm. Each peak fraction collected was vacuum-dried (Savant Instrument Inc^®^), reconstituted in 1 mL ultrapure water and evaluated by antimicrobial activity assays.

### 2.4 MICROBIAL STRAINS

Bacterial and fungal strains were obtained from the collection of microorganisms of the Laboratory for Applied Toxinology (LETA) of the Butantan Institute (São Paulo, Brazil). The bioassays were performed with Gram-positive bacteria *Bacillus megaterium* ATCC10778, *Bacillus subtilis* ATCC6633, *Micrococcus luteus* A270, *Staphylococcus aureus* ATCC29213, Gram-negative Bacteria *Enterobacter cloacae* β-12, *Escherichia coli* SBS 363, *Salmonella arizonae* ATCC13314, Filamentous Fungi *Aspergillus niger* (isolated bread), *Beauveria bassiana* (isolated from mummified insect), *Cladosporium sp* (isolated bread) and Yeast *Candida glabrata* IOC4565, *Candida krusei* IOC4559 and *Cryptococcus neoformans var. neoformans* B-350 1A.

### 2.5 ANTIMICROBIAL ASSAYS

The antimicrobial effects of the fraction were evaluated by liquid growth inhibition assays as Riciluca [25], using *Bacillus megaterium* ATCC10778, *Bacillus subtilis* ATCC6633, *Micrococcus luteus* A270, *Staphylococcus aureus* ATCC29213, *Enterobacter cloacae* β-12, *Escherichia coli* SBS 363, *Salmonella arizonae* ATCC13314, *Aspergillus niger* (isolated bread), *Beauveria bassiana* (isolated from mummified insect), *Cladosporium sp* (isolated bread), *Candida glabrata* IOC4565, *Candida krusei* IOC4559 and *Cryptococcus neoformans var. neoformans* B-350 1A. Bacteria were cultured in poor nutrient broth (PB) (1.0 g peptone in 100 ml of water containing 86 mM NaCl at pH 7.4; 217 mOsM) and fungus and yeasts were cultured in poor potato dextrose broth (1/2 PDB: 1.2 g potato dextrose in 100 ml of H_2_O at pH 5.0; 79 mOsM). Determination of antimicrobial peptide was performed using 5-fold microtiter broth dilution assay in 96-well sterile plates at a final volume of 100 μL. Mid-log phase culture was diluted to a final concentration of 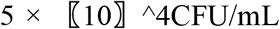 for bacteria and 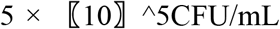 for fungus and yeast, as Segura-Ramírez & Silva Júnior [26]. Dried fractions were dissolved in 200 μL of ultrapure water and 20 μL aliquoted into each well with 80 μL of the microbial dilution. The assays were executed in duplicate. The microplates were incubated for 18 h at 30 °C, under constant agitation. Growth inhibition was determined by measuring absorbance at 595 nm.

### 2.6 HEMOLYTIC ASSAY

Human erythrocytes from a healthy adult donor were collected with 0.15 M citrate buffer and washed three times by centrifugation (800 x g, 15 min, 4 °C), the supernatant was discarded. Aliquots of 100 μL in a 3% (v/v) suspension of washed erythrocytes in 0.15 M phosphate-buffered saline (PBS) was incubated with the fraction in a U-bottom 96-well microplate for 3 h at 37 °C with constant shaking. The supernatant was transferred to a 96-well flat microplate and the hemolysis was measured by the absorbance at 405 nm of each well in a microplate reader Victor^3^ (1420 Multilabel Counter/Victor3, Perkin Elmer). The hemolysis percentage was expressed in relation to a 100% lysis control (erythrocytes incubated with 0.1% triton X-100); PBS was used as a negative control. The equation used for % of hemolysis was: % Hemolysis: [(Abs pep – Abs PBS)/Abs Triton 0.1% - Abs PBS) *100]

### 2.7 MASS SPECTROMETRY AND BIOINFORMATICS ANALYSIS

Samples were suspended in 15 μL of 0.1% formic acid solution and analyzed by liquid chromatography coupled to tandem mass spectrometry (LC-MS/MS) on Thermo Scientific™ LTQ XL^™^ - ETD mass spectrometer (Thermo Fisher Scientific, Bremen, Germany) coupled to an Easy-nLC 1000 (Thermo Fisher Scientific, Bremen, Germany). The samples were (10μL) automatically injected into a Júpiter C18 (10 μm, 100 μm x 50 mm) pre-column (Phenomenex) coupled to a C18 capillary analytical reverse phase column. A linear gradient of 5 to 95% acetonitrile/formic acid 0,1% for 30 minutes and a flow rate of 140 nL / min was used. The ionization source was operated in positive mode, which detects positively charged ions. The spectra were collected and analyzed in Xcalibur 2.0 software (Thermo Electron, USA). The deconvolution of the m / z values to obtain the molecular weight of the protein was performed in the software MassLynx V4.1 Walters^®^ (accessed on 20th August 2020) and PEAKS^®^X+ Studio software (v10.5; Bioinformatics Solutions, Waterloo, ON, Canada). Also, peptides were searched against the NCBI database using the PEAKS DB tool with Gastropoda, *Limax* and *Limacus flavus* databases and compared to Li et al. [24], KEGG pathways. The physicochemical parameters were evaluated in the bioinformatics tool ChemDraw and Chem3D Professional 16.0.4 and with the online programs as PepCalc (https://pepcalc.com/), PepDraw (http://pepdraw.com/), all accessed on 22th August 2020 and ProteinBlast (https://blast.ncbi.nlm.nih.gov) tool.

## 3. RESULTS

### 3.1 FRACTIONATION OF THE MUCUS AND ANTIMICROBIAL SCREENING

The mucus extract from three adult specimens was processed as previously described. The resulting supernatant was applied to a reverse-phase HPLC and was isolated 3 fractions (Figure 2) with antimicrobial activity when analyzed in liquid growth inhibitory assays. All fractions were tested against 13 microorganisms’ strains and showed some type of antimicrobial activity (Table 1). Fractions were named as LFMP-Fp001, LFMP-Fp002 and LFMP-Fp003. Fractions, LFMP-Fp001 and LFMP-Fp002, inhibited only *B. subtillis* and *A. niger* however LFMP-Fp003 showed greater activities, inhibiting *B. subtillis, C. neoformans, E. cloacae* and *M. luteus* A270.

**Figure 2.**
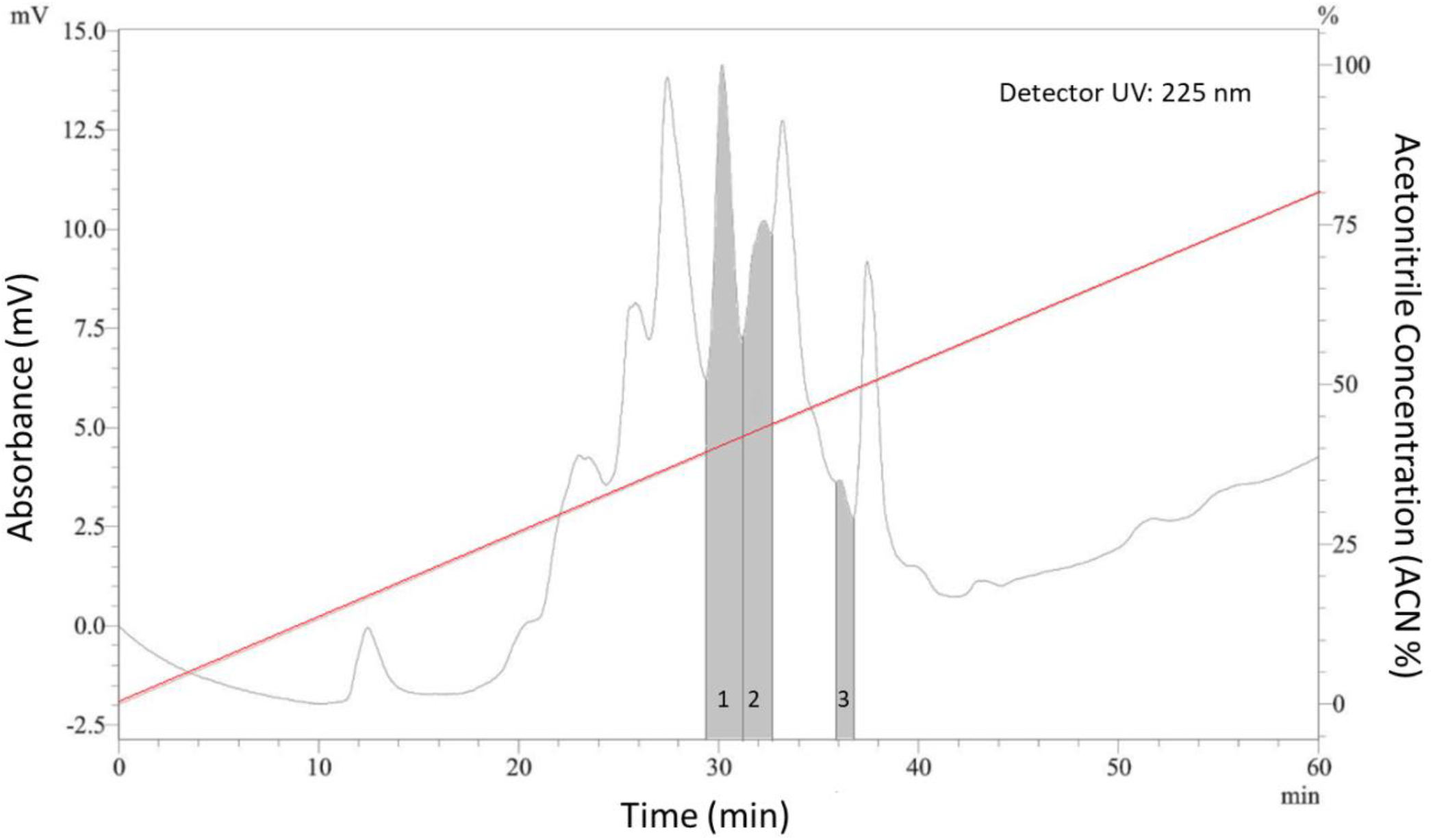
Fractionation of the *Limacus flavus* mucus extract in 10% DMSO (eluted in 80% acetonitrile). Chromatographic fractionation profile by high performance liquid chromatography in reversed-phase using Shim-pack XR-ODS C18 preparative column of mucus treated with 10% DMSO, with flow 8.0 mL / min, in a linear gradient from 0% to 80% of ACN at 0.01% TFA in 60 min; 225 nm absorbance.The enumerated peaks correspond to the fractions with antimicrobial activity.

**Table 1.**
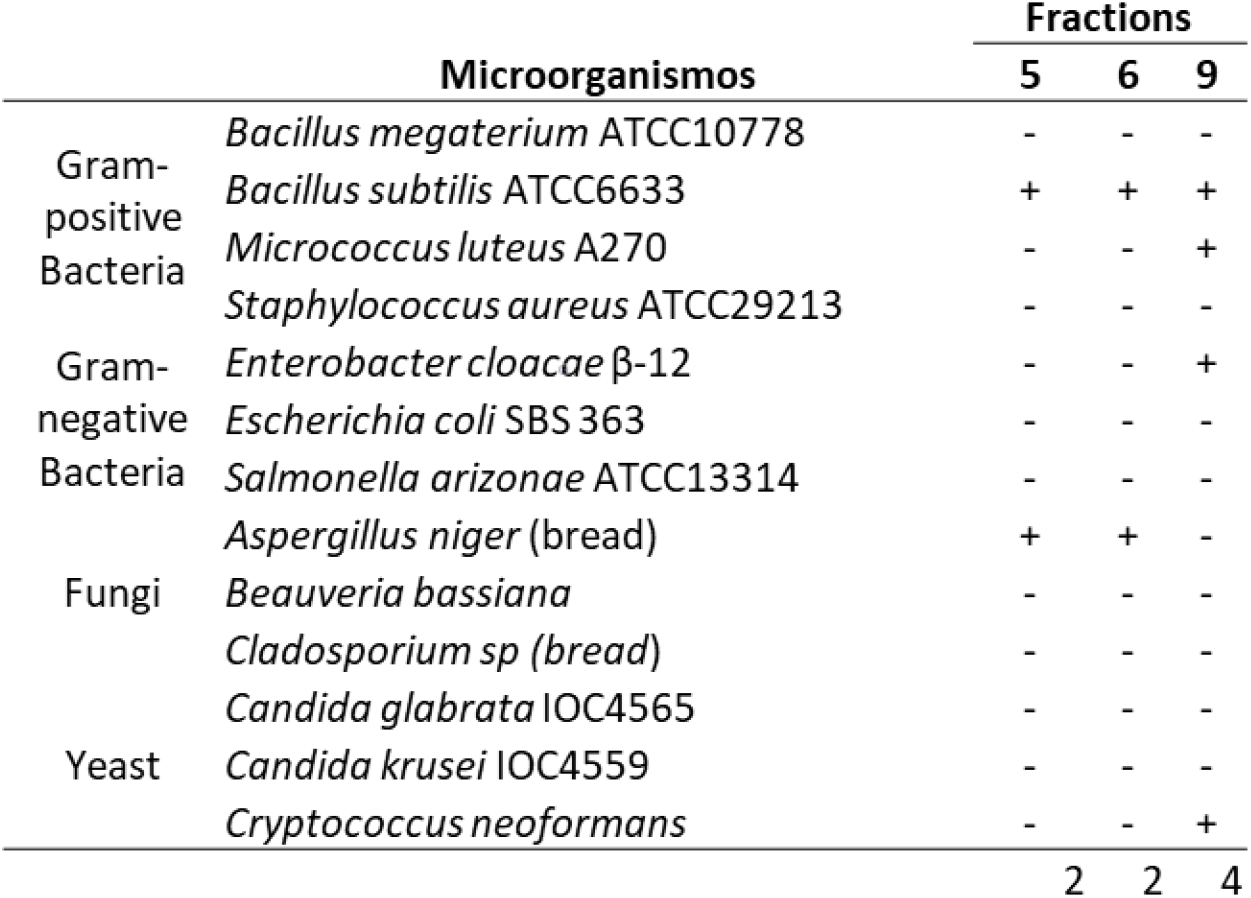
Antimicrobial activity in liquid growth of *Limacus flavus* slug mucus fractions collected by high-performance liquid chromatography in reversed-phase using Shim-pack XR-ODS C18 analytical column of mucus treated with 10% DMSO, with flow 2.0 mL / min, in 60 min; 225 nm absorbance. Tests carried out against *Aspergillus niger* (bread isolate*), Beauveria bassiana, Bacillus megaterium* ATCC10778, *Bacillus subtilis* ATCC6633, *Candida glabrata* IOC4565, *Candida krusei* IOC4559, *Cladosporium sp* (bread isolate)*, Cryptococcus neoformans, Enterobacter cloacae* 363, *Micrococcus luteus* A270, *Salmonella arizonae* ATCC13314 and *Staphylococcus aureus* ATCC29213. The symbols: (+) inhibition of antimicrobial activity and (-) non-inhibition of antimicrobial activity.

### 3.2 HEMOLYTIC ASSAY

All fractions were tested for hemolytic assay, no hemoglobin release was observed. The three fractions show non-toxicity activity against human erythrocytes.

### 3.3 MASS SPECTROMETRY AND BIOINFORMATICS ANALYSIS

By the deconvolution of the fractions by PeaksX+ software, all fractions presented mass less than 3 kDa. Characterization of LFMP-Fp001 primary structure by MS/MS exhibited 636.74 Da with sequence YLRW (Figure 3); LFMP-Fp002, 560.6 Da sequence DLQW (Figure 4) and LFMP-Fp003, 560.6 Da with sequence WLGH (Figure 5). When LFMP-Fp001 was processed in NCBI database (Blastp) (Table 2) LFMP-FP001 showed similarities to *Aplysia californica, Biophalaria glabrata, Lottia gigantea* and *Pomacea canaliculata* proteins. When compared to KEGG pathways (Table 3), none of the proteins listed are related to immune system pathways. When compared to NCBI database (Table 4) the sequence of LFMP-Fp002 matched with 6 organisms, *A. californica, B. glabrata, Elysia chlorotica, Littorina littorea, Lottia gigantean* and *P. canaliculata*. Most of the proteins found are isoforms or uncharacterized/hypothetical sequences. When compared to KEGG pathways, *L. littorea* presents two toll-like receptors proteins. In addition, *P. canaliculata* exhibited similarities with CD109 antigen-like proteins and Mitogen-activated protein kinase 13-like.The fraction LFMP-FP003 showed similarities to the same organisms as the fraction LFMP-FP001, being *B. glabrata* and *P. canaliculata* with most proteins identifications (Table 5). Presenting the same characteristics, most of the sequences are not identified and not related to KEGG pathways.

**Figure 3.**
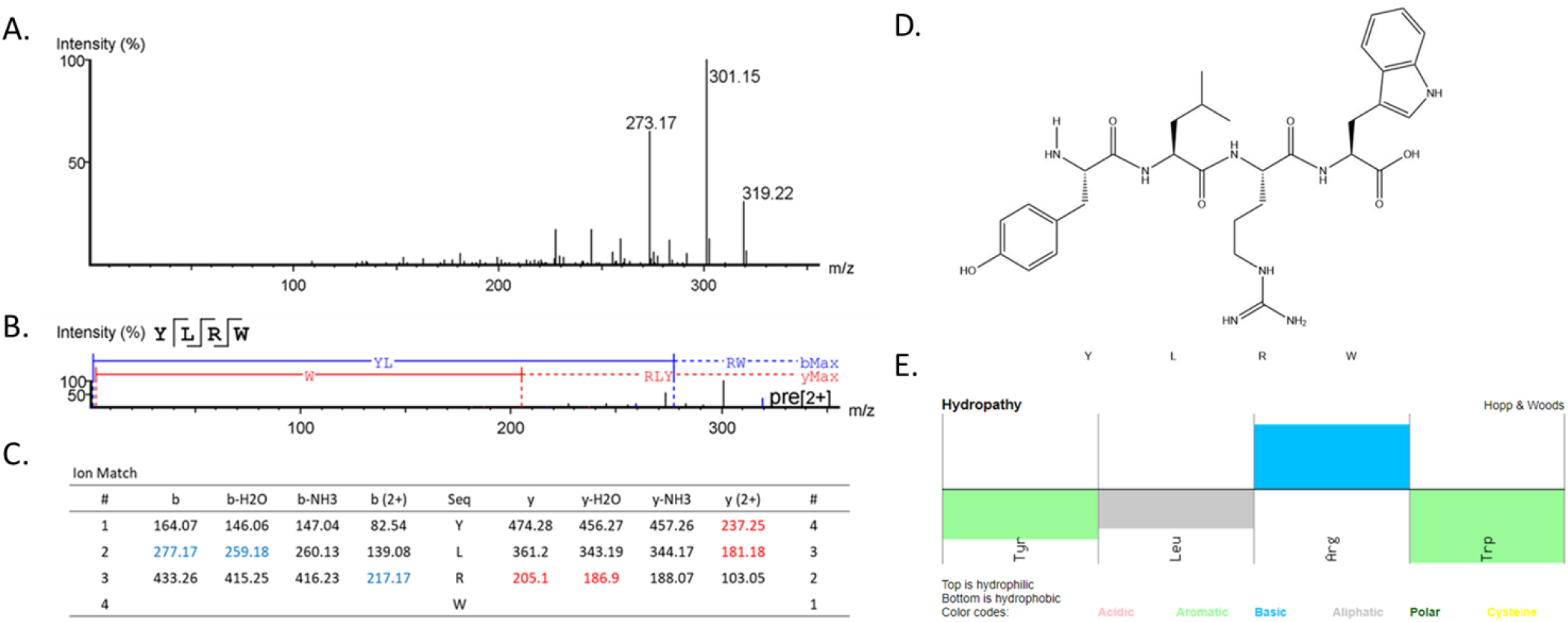
Deconvolution profile of the LFMP-Fp001 fraction after mass spectrometry (ESI-Q-Tof / MS) on Thermo Scientific™ LTQ XL™ - ETD mass spectrometer coupled to an Easy-nLC 1000 (A) Profile of the mass spectrometer using bioinformatic Peaks^®^X+ tool (B) Collision-induced dissociation (CID) spectrum of the de novo sequenced peptide. The ions relative to -y (red) and -b (blue) series indicated in the spectrum correspond to the amino acid sequence of the antimicrobial peptide, YLRW. The sequence is represented by standard amino acid code letters. (C) Ion matches profile of YLRW (D) Primary structure of YLRW accessed by the PerkinElmer ChemDraw Professional tool (E) Hydropathy parameters of LFMP-Fp001 by PepCalc tool.

**Figure 4.**
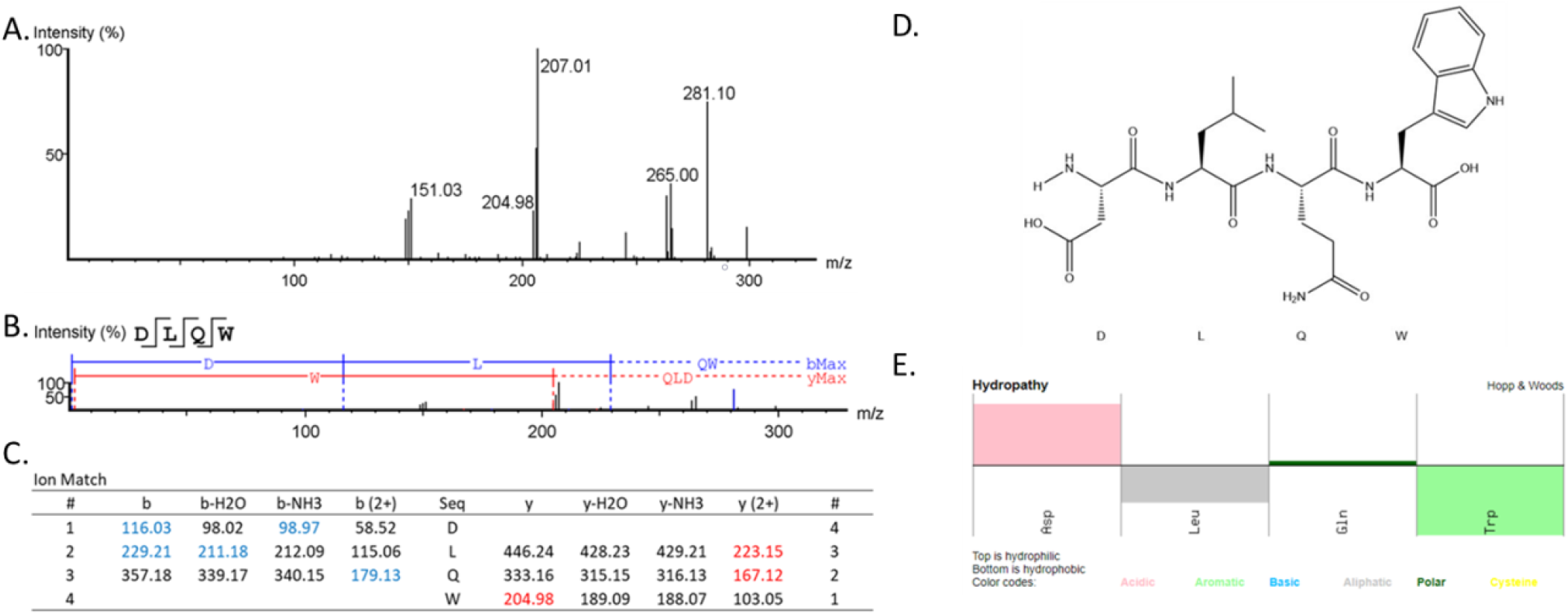
Deconvolution profile of the LFMP-Fp002 fraction after mass spectrometry (ESI-Q-Tof / MS) on Thermo Scientific™ LTQ XL™ - ETD mass spectrometer coupled to an Easy-nLC 1000 (A) Profile of the mass spectrometer using bioinformatic Peaks^®^X+ tool (B) Collision-induced dissociation (CID) spectrum of the de novo sequenced peptide. The ions relative to -y (red) and -b (blue) series indicated in the spectrum correspond to the amino acid sequence of the antimicrobial peptide, DLQW. The sequence is represented by standard amino acid code letters. (C) Ion matches profile of DLQW (D) Primary structure of YLRW accessed by the PerkinElmer ChemDraw Professional tool (E) Hydropathy parameters of LFMP-Fp002 by PepCalc tool.

**Figure 5.**
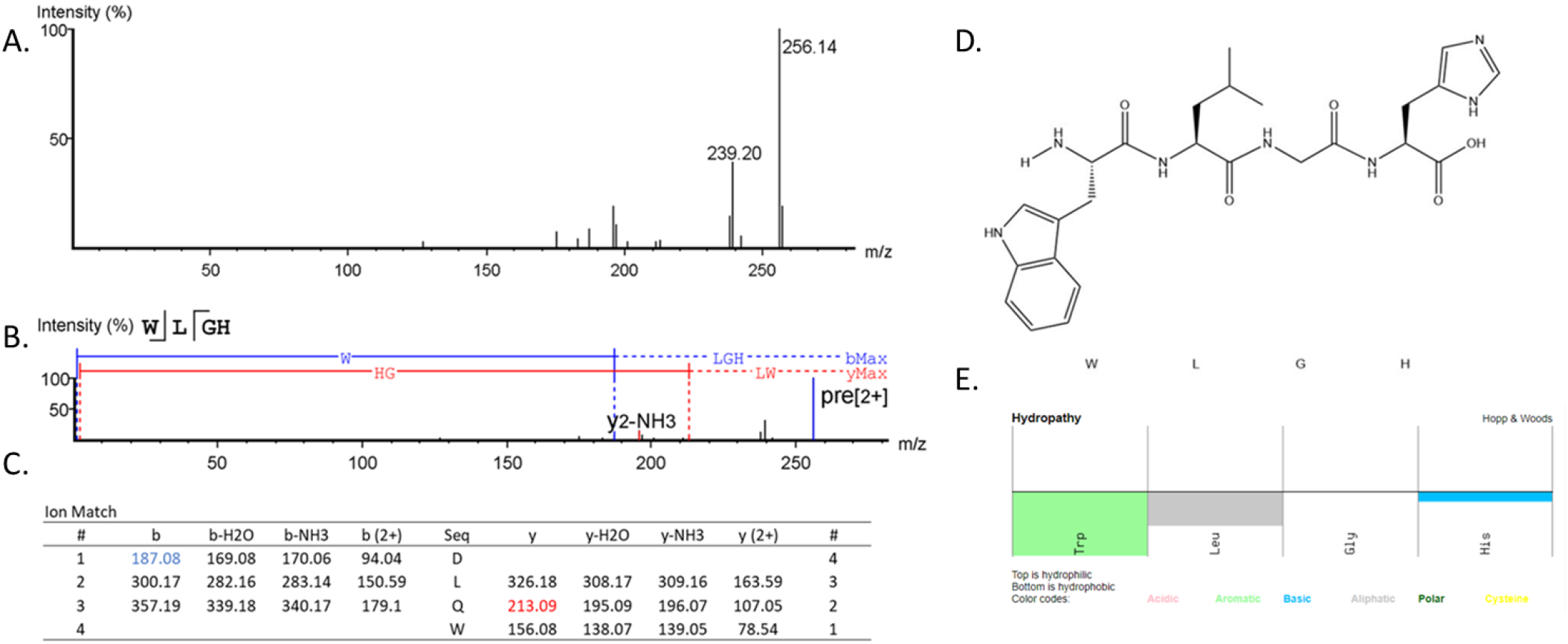
Deconvolution profile of the LFMP-Fp003 fraction after mass spectrometry (ESI-Q-Tof / MS) on Thermo Scientific™ LTQ XL™ - ETD mass spectrometer coupled to an Easy-nLC 1000 (A) Profile of the mass spectrometer using bioinformatic Peaks^®^X+ tool (B) Collision-induced dissociation (CID) spectrum of the de novo sequenced peptide. The ions relative to -y (red) and -b (blue) series indicated in the spectrum correspond to the amino acid sequence of the antimicrobial peptide, WLGH. The sequence is represented by standard amino acid code letters. (C) Ion matches profile of WLGH (D) Primary structure of YLRW accessed by the PerkinElmer ChemDraw Professional tool (E) Hydropathy parameters of LFMP-Fp003 by PepCalc tool.

**Table 2.**
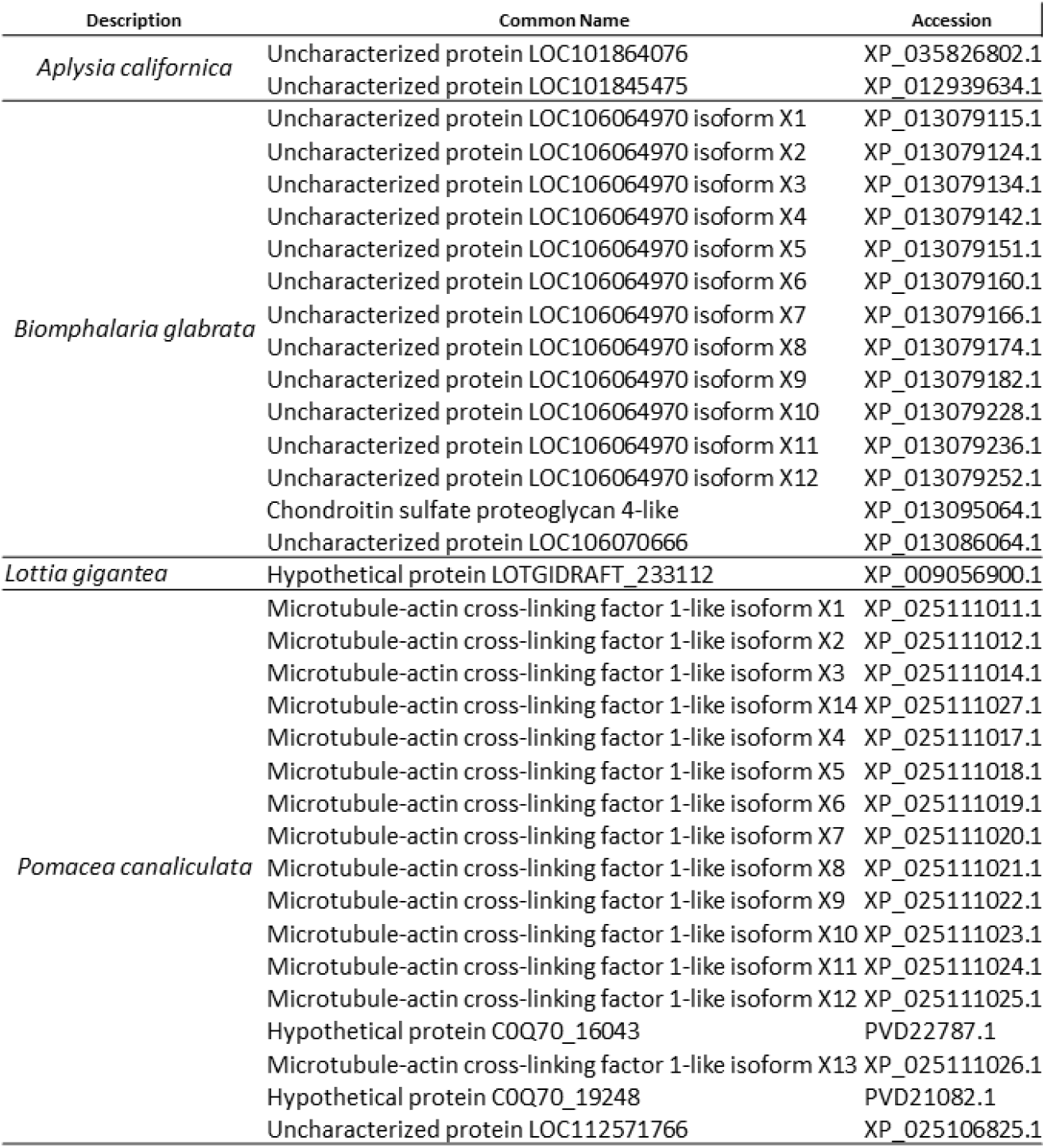
Aligments results of the fraction LFMP-Fp001 with ProteinBlast (blastp suite) from NCBI. Results with E value of 49, Total score of 19.3, Query cover of 100% and restrict to Gastropoda (taxi: 6448), accessed in 11th December 2020.

**Table 3.** KEGG analysis of the immune system pathways from the *Limacus flavus* body’s transcriptome of Li et al., 2020.

| KEGG Pathway | Pathway ID | Gene Number |
| --- | --- | --- |
| Antigen processing and presentation | ko04612 | 128 |
| B cell receptor signaling pathway | ko04662 | 73 |
| Chemokine signaling pathway | ko04062 | 149 |
| Complement and coagulation cascades | ko04610 | 24 |
| Cytosolic DNA-sensing pathway | ko04623 | 42 |
| Fc epsilon RI signaling pathway | ko04664 | 71 |
| Fc gamma R-mediated phagocytosis | ko04666 | 120 |
| Hematopoietic cell lineage | ko04640 | 18 |
| Intestinal immune network for IgA production | ko04672 | 1 |
| Leukocyte transendothelial migration | ko04670 | 135 |
| NOD-like receptor signaling pathway | ko04621 | 60 |
| Natural killer cell mediated cytotoxicity | ko04650 | 69 |
| Platelet activation | ko04611 | 185 |
| RIG-I-like receptor signaling pathway | ko04622 | 37 |
| T cell receptor signaling pathway | ko04660 | 94 |
| Toll-like receptor signaling pathway | ko04620 | 68 |

**Table 4.**
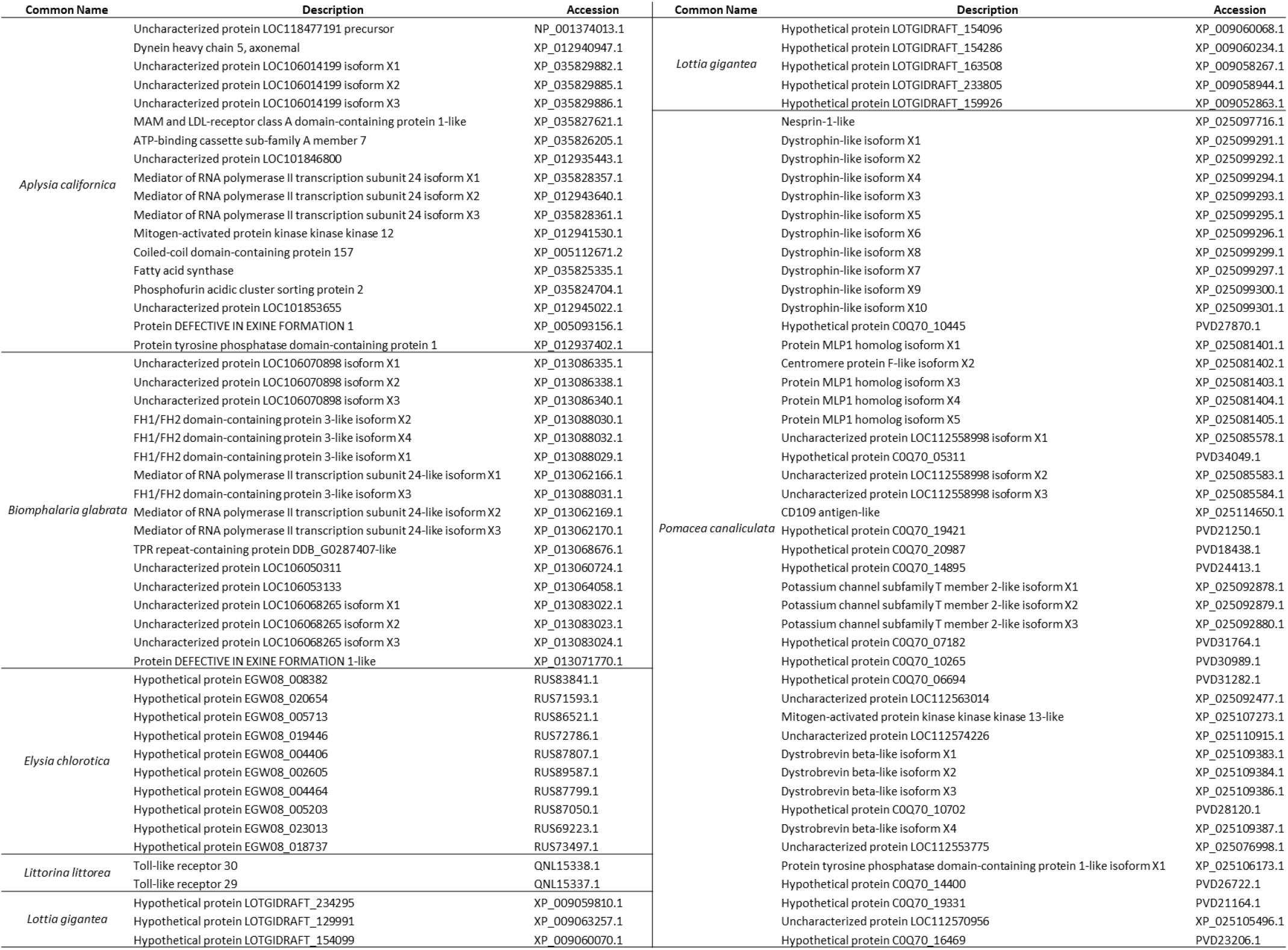
Aligments results of the fraction LFMP-Fp002 with ProteinBlast (blastp suite) from NCBI. Results with E value of 99, Total score of 33.9, Query cover of 100% and restrict to Gastropoda (taxi: 6448), accessed in 11th December 2020.

**Table 5.**
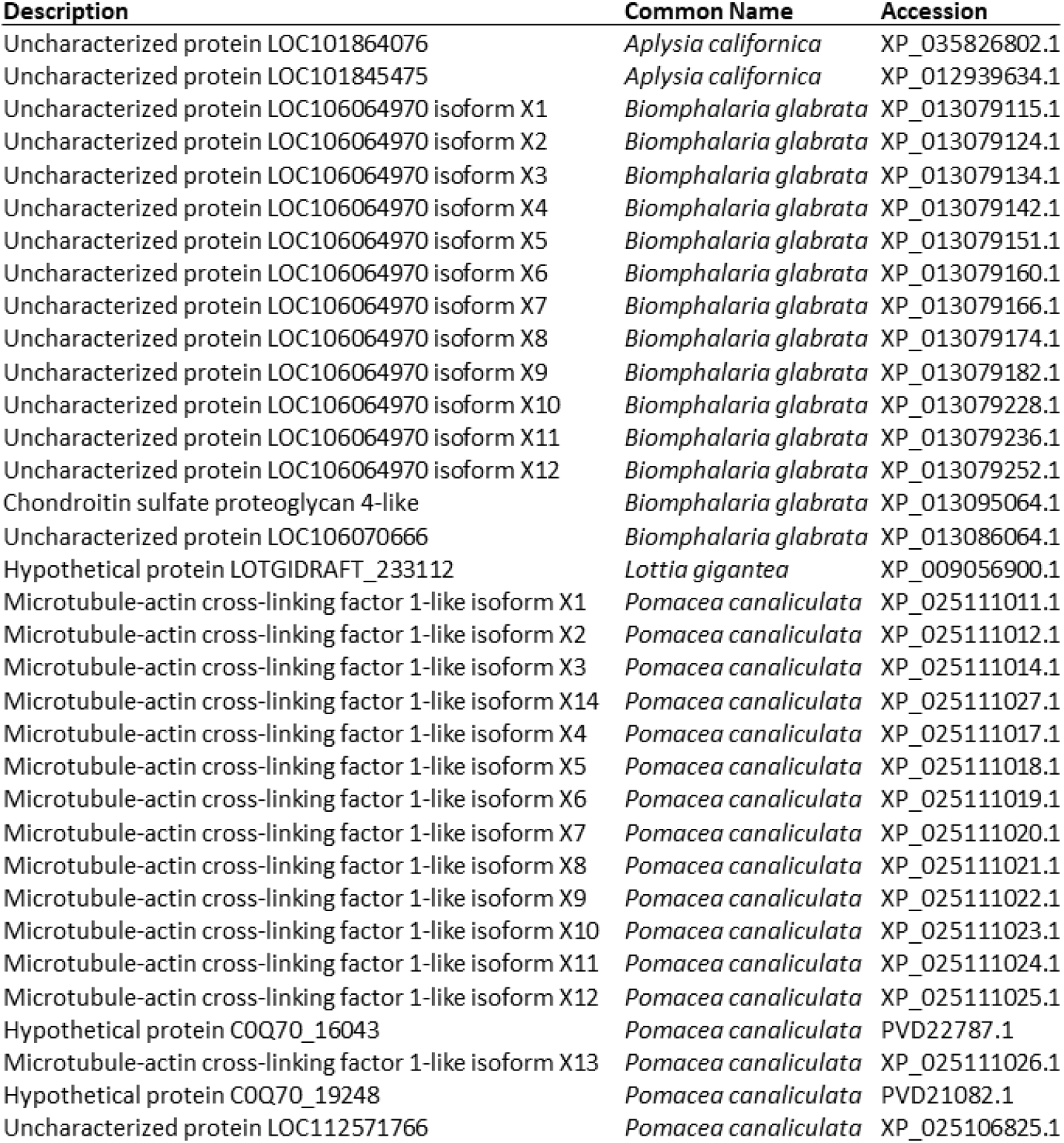
Aligments results of the fraction LFMP-Fp003 with ProteinBlast (blastp suite) from NCBI. Results with E value of 140, Total score of 18.0, Query cover of 100% and restrict to Gastropoda (taxi: 6448), accessed in 11th December 2020.

### 3.4 STRUCTURE AND PHYSIC-CHEMICAL CHARACTERIZATION

Sequences similarity searches with Basic Local Alignment Search Tool (BLAST) were performed but there was no significant similarity found for the three fractions, due to its short sequence predicted. Physical and chemical characteristics were predicted by bioinformatics online programs which net charge, theoretical isoelectric point (pI) and solubility were predicted. LFMP-Fp001 is a cationic molecule (net charge of 1) due to the presence in its structure of one tyrosine (Y), one leucine (L), one positively charged arginine residue (R) and one tryptophan (W). Furthermore the peptide has a pI of 9.57, indicating that this is the pH value at which its net charge is equal to 0. By estimating its solubility, LFMP-Fp001 has poor water solubility, considerate a hydrophobic AMP. LFMP-Fp002 is an anionic peptide (net charge of - 1) due to one negatively charged aspartic acid (D), one leucine (L), one glutamine (Q) and one tryptophan (W) and presents an iso-electric point of pH 0.67. Based on the iso-electric point, the number of charged residues, and the peptide length of four residues, this peptide might have good water solubility, an hydrophilic AMP. LFMP-Fp003 is a neutral peptide (net charge zero) due to its sequence, as one tryptophan (W), one leucine (L), one glycine (G) and one histidine (H), presents an iso-electric point of pH 7.69 and has poor water solubility, being considerate an hydrophobic AMP.

## 4. DISCUSSION

One of the main crises in the public health system worldwide is the antimicrobial resistance (AMR) [1] and since the Antibiotic Revolution, natural products from a wilde range of organism are being studied for suppress diseases and deaths [27]. The raw mucus presented an elastic consistency and translucent-yellowish color, same described by Boffi [28] and Barker [29] and after liophilization it presented a cotton-like texture. Thermal shock method was chosen for being the less aggressive treatment to collect more material of the species, as described by Pemberton [30], otherwise saline solution could extract unlike residues. Among the organisms tested, it is noted that the mucus showed greater inhibition for filamentous fungi and Gram-positive bacteria, followed by Gram-negative bacteria and fungi yeasts. The assay established sensitivity to a range of microorganisms which can be related as probable pathogens that the Phylum Mollusca can be exposed in the environments [31]. The Phylum Mollusca is one of the most diverse phyla in the animal kingdom; they originated from the Cambrian period [32] and have undergone few changes in the course of evolution [33]. Its irradiation allowed colonization in various environments (ocean, fresh water and land) [34], enriched with pathogens, such as viruses, fungi and bacteria. Currently it is estimated that mucus presented antimicrobial properties beyond the breathing and hydration functions, communication, locomotion and adhesion in surfaces [19]. All the fractions were tested for hemolytic assay, showing. These data suggest that the mechanism of action of the *L. flavus* mucus does not involve the disruption of cell membranes. Once you consider developing novel antibiotics for human application, the drug itself must show low toxicity against erythrocytes [35], which all three fractions demonstrated.

When analyzed by mass spectrometry and bioinformatics tools, LFMP-Fp001 did not show similarities with KEGG pathways of Li et al.[24], instead when the BLAST was proceed, it showed similarities with close organisms although most of the proteins were uncharacterized/hypothetical. Comparing the results from BLAST and KEGG pathways, LFMP-Fp002 can be related to two proteins matches: *L. littorea* presents two toll-like receptors proteins and *P. canaliculata*, CD109 antigen-like and Mitogen-activated protein kinase 13-like. Toll-like receptor signaling pathway (KO04620), which recognizes pathogen-associated molecular patterns derived from microbes. CD109 antigen is related to T-cell, once these cells receptors are activated, it initiates the signaling process of cascades to respond to the infection; and Mitogen-activated protein in humans is an important role in the cascades of cellular responses evoked by extracellular stimuli such as pro-inflammatory cytokines or physical stress. Once applied these proteins like to *L. flavus* slugs, they may have a similar processes, since these organism is constantly exposed to a wide range of microbes and external stresses. The fraction LFMP-FP003 faced the same situation as LFMP-FP001, although it presented similarities with some invertebrates, none of the proteins were characterized, same to proteins identified to KEGG pathways. The three fractions had different physic-chemical characteristics and when thought of a novel antibiotic, these characteristics are important when it comes to the variety of microorganisms, target and biological responses, since AMR is of global health problem.

## 5. CONCLUSION

In conclusion, we isolated, fractionated and characterized the three antimicrobial peptides of the mucus of yellow slug *L. flavus*. The three peptides LFMP-FP001, LFMP-FP002 and LFMP-FP003 presents a wide range of antimicrobial activity, showing more sensibility to Gram-positive bacteria and filamentous fungi, following to Gram-negative bacteria and fungi yeast. Physic-chemical characteristics were elucidated, which LFMP-FP001 is featured as a cationic AMP, LFMP-FP002 an anionic AMP and LFMP-FP003 neutral AMP (no net charge). Furthermore, none of the molecules has a cytotoxic activity against human erythrocytes, suggesting that the three AMPs have potential for a novel antibiotic. Thus, further investigations will be carried out on multidrug resistant bacteria and mode of action of the three molecules, considering antimicrobial resistance is one of the major public health problems. Finally, it is worth highlight that this work complements the literature of *L. flavus* and indicates that the mucus of this species can be a source of antimicrobial molecules.

## ETHICS APPROVAL AND CONSENT TO PARTICIPATE

This research was approved and performed in accordance with the Ethical Principles in Animal Research adopted by the Ethics Committee in the Use of Animals of Butantan Institute (N° 5649250717) and Plataforma Brasil CAAE N°19403819.1.0000

## ACKNOWLEDGEMENTS

Conceptualization, P.Y.H. and P.I.S.J.; Data curation, P.Y.H.; Formal analysis, P.I.S.J.; Funding acquisition, P.I.S.J.; Investigation, P.Y.H.; Methodology, P.I.S.J.; Project administration, P.I.S.J.; Resources, P.I.S.J.; Supervision, P.I.S.J.; Validation, P.Y.H. and P.I.S.J.; Writing-original draft, P.Y.H.; Writing-review and editing, P.Y.H. and P.I.S.J.

Special thanks to Dr. Milton Yutaka Nishiyama Junior (Center for Bioinformatics and Computational Biology, Butantan Institute, São Paulo (SP), Brazil) for helping us with the databases.

